# Classification of iPSC-Derived Cultures Using Convolutional Neural Networks to Identify Single Differentiated Neurons for Isolation or Measurement

**DOI:** 10.1101/2023.12.24.573194

**Authors:** Purva Patel, Lina K Mohammed Ali, Uma Kaushik, Mallory G Wright, Kaylee P Green, Jason E Waligorski, Colin L Kremitzki, Graham W Bachman, Serena N Elia, William J Buchser

## Abstract

Understanding neurodegenerative disease pathology depends on a close examination of neurons and their processes. However, image-based single-cell analyses of neurons often require laborious and time-consuming manual classification tasks. Here, we present a machine learning approach leveraging convolutional neural network (CNN) classifiers that have the capability to accurately identify various classes of neuronal images, including single neurons. We developed the Single Neuron Identification Model 20-Class (SNIM20) which was trained on a dataset of induced pluripotent stem cell (iPSC)-derived motor neurons, containing over 12,000 images from 20 distinct classes. SNIM20 is built in TensorFlow and trained on images of differentiated iPSC cultures stained for nuclei and microtubules. This classifier demonstrated high predictive accuracy (AUC = 0.99) for distinguishing single neurons. Additionally, the 2-stage training framework can be used more broadly for cellular classification tasks. A variation was successfully trained on images of a human osteosarcoma cell line (U2OS) for single-cell classification (AUC = 0.99). While this framework was primarily designed for single-cell microraft-based identification and capture, it also works with cells in standard plate formats. We additionally explore the impact of specific fluorescent channels and brightfield images, class groupings, and transfer learning on the quality of the classification. This framework can both assist in high throughput neuronal or cellular identification and be used to train a custom classifier for the user’s needs.

## Introduction

Research on neurodegenerative diseases such as ALS, neuropathies, and Parkinson’s Disease centers on the study of neurons, investigating the damage to these cells and their molecular causes. Studying single-cell neuronal phenotypes often necessitates meticulous efforts in manual cellular annotation due to highly entangled neurites, which present a challenge in connecting an axon to its corresponding cell body. For massively parallel approaches to study the impact of genetic variants on neuronal morphology or organellar phenotypes, a pooled assay is used, which requires single-cell evaluation and nuclear identification or capture. Cultures of induced pluripotent stem cell (iPSC)-derived neurons on microrafts allows single neurons to be discretized for phenotyping and single-cell isolation^1^. Each raft (akin to a region of interest that can be captured) can be annotated to determine its contents, such as a single differentiated neuron, an undifferentiated cell, a dead cell, etc. Here, we outline a pipeline and present pretrained machine learning (ML) models that enable rapid classification and automatically annotate regions of microscopic images.

The use of artificial neural networks (aNNs) to classify cells has become simpler due to the availability of manually annotated datasets containing biological and medical images, and platforms like the Allen Cell Explorer^2,3^. However, it remains a challenge to identify microscopic images of individual neurons, since many cultures or slices contain a mixture of different cell types (neurons, undifferentiated cells, dead cells), and the healthy neurons form networks and clusters which facilitate neuronal growth but make segmentation difficult. While there are image-based algorithms and ML tools that enable neuronal segmentation, cell tracing, and identification of neurons, *single* neuron analyses remain difficult ^4–9^. aNNs make an ideal choice to overcome limitations in the existing tools and have been implemented in similar contexts^10–15^.

Here, we use and share a set of single iPSC-derived human motor neuron images and aNNs to make a neuronal classifier and present a pipeline to quickly train other custom aNNs for regional annotations. We built “single neuron imaging model 20-class” (SNIM20), a convolutional neural network (CNN) that automates manual annotation of images in a high throughput setting. SNIM20 can identify and classify images of single iPSC-derived motor neurons with high accuracy, which can then be taken into a variety of downstream applications, such as organelle quantification in the distal neurite. We conducted a comparative analysis of the SNIM20 classifier against a variety of related classifiers that varied in their fluorescent wavelengths, class composition, and model architecture. We also applied a similar CNN-based architecture to a non-neuronal cell type to evaluate the model’s performance and versatility across different experimental paradigms. This approach highlights the adaptable nature of image CNN classifiers for annotating various types of cellular images.

## Results

### Overview of the Imaging and Training Pipeline

To enable pooled, image-based functional genomics pipelines, we sought to image motor neurons in a high-throughput manner. We differentiated iPS cells into motor neurons (**Fig. S1**) and then seeded them onto a microraft surface. We imaged the differentiated neuronal cultures to visualize the nucleus and the tubulin cytoskeleton (**Fig. 1a**). The training images are then hand-annotated locally using FIVTools (in-house developed software which enables multi team-member annotation and image export) (**Fig. 1b, Fig. S2**). The name of the folders that the images are placed in designates their class (which is dependent on the experiment). We independently trained several CNN classifiers, then selected the best one, therefore giving rise to an initial model (designated as v1 in **Fig. 1a**). The second training round starts by using the ‘v1’ classifier to automatically annotate the classes on a new set of images. These inferred images are then exported if they have a relatively high prediction score. Since the ‘v1’ classifier has had minimal training, it can make mistakes, necessitating some manual re-sorting prior to a new round of training and selecting a new ‘v2’ classifier which is much more robust (**Fig. 1c**). This process allows the initial training set to be relatively small, then the machine learning can ‘assist’ to get a much larger training set which is used for subsequent training, resulting in a highly tuned classifier.

**Figure 1:**
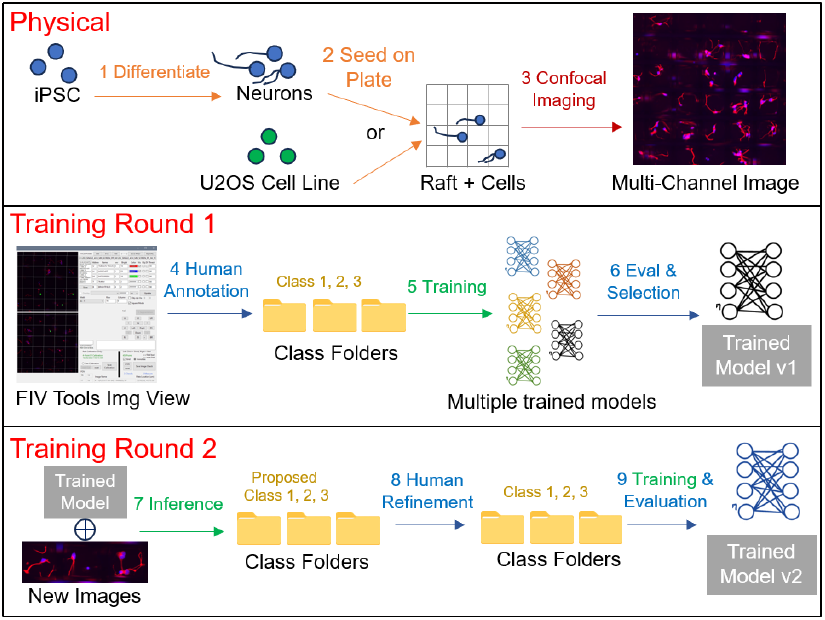
Schematic representation of neuronal identification pipeline. **a**. Physical. Schematic representation of automated class annotation workflow. For figures in the manuscript, this either involves differentiated motor neurons or U2OS cancer cell line seeded onto microraft plates, but also works in a standard plate format. Cells are stained and later imaged using fluorescent microscopy. **b**. Training Round 1. In-house custom software, FIVTools, was used to manually annotated and then export images into different folders of interest, named with the class label. Using this dataset, we trained many initial convolutional neural network (CNN) classifiers. These were automatically evaluated, and the best initial model selected as the ‘v1’ classifier. **c**. Training Round 2. Upon acquisition of additional training data, images were automatically scored and sorted into their labeled class folder by the ‘v1’ classifier. The images in these folders are then manually reviewed and re-sorted as needed. The second round of training then utilizes the combination of the original training set and this larger set of model-assisted annotations to further refine and train a new iteration of the classifier, designated ‘v2’.

### Differentiated Neurons and Class Definitions

Experiments with differentiated neurons on rafts led to many individual outcomes (**Fig. 2a** and **Fig. S3**). We broadly classified the training set images into four groupings: “No nuclei,” “One nuclei,” “Two nuclei,” and “Multiple nuclei” (three or more). We were focused on identifying high-quality single and double neurons with distinct nuclei and neurites. These groupings guided our selection of rafts for isolation and downstream genotyping, owing to our interest in single-cell phenotypic analyses. However, we observed variations within the “one nuclei” grouping, particularly neurons with a very short neurite, disconnected neurite, and those with poor focus, so each of those were made into a distinct class. We also introduced multiple classes within the “Two nuclei” grouping (**Fig. 2b, Fig. S4**). In total, there were 12,689 images in the training set (**Table S1**). Just as for human annotators, the classifiers had more difficulty distinguishing between classes like “1 Cell and Disconnected Neurite,” “1 Neuron,” “1 Neuron Poor Focus,” and “2 Nuclei 1 Soma” (**Fig. 2c**). Confusion between one and two neurons per raft can lead to errors in interpretation of downstream molecular results derived from sequencing. Figure 2b also acted like a ‘legend’ and these classes were used to define the “SNIM20” classifier. After training, SNIM20 consisted of varying filter sizes within its convolutional layers to capture diverse image features at different scales (**Fig. 2d**). While a different biological question would have different classes, the details of the class types are presented to show the possible subtlety that can be captured by these classifiers. Similarly, the architecture shown is the one that worked the best, but additional optimization or a different training set would result in variations of the parameters and architecture that are likely to work just as well.

**Figure 2:**
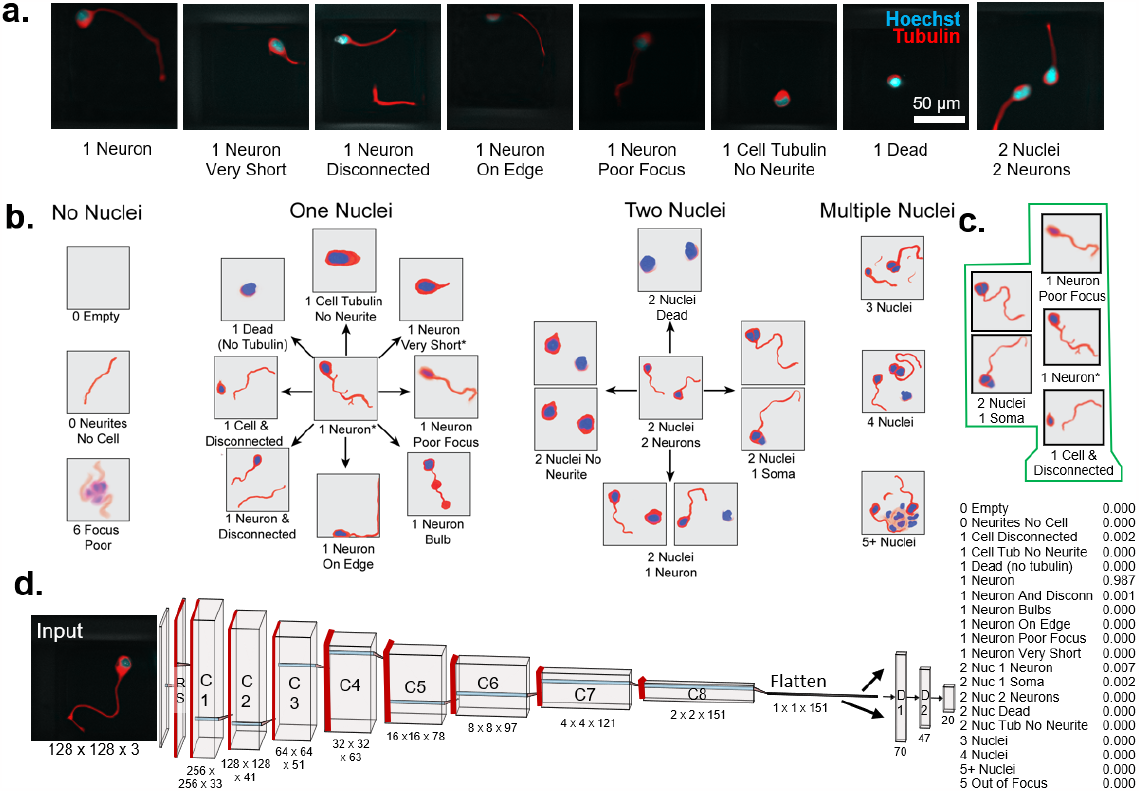
Classification schema for differentiated neurons on microrafts. **a**. Confocal microscope images for select classes in the training set for the SNIM20 classifier. Motor neuron cultures were live stained and imaged to visualize the tubulin cytoskeleton (red), and nuclei (blue). **b**. Schematic of the four class groupings, including rafts that have no nuclei, one nuclei, two nuclei, and three or more nuclei. Within each grouping, cartoon depictions of the respective classes are shown. The asterisk indicates the desired neuronal phenotype in these experiments (true single neurons). Within the two nuclei grouping, there are two closely related examples for “2 Nuclei No Neurite,” “2 Nuclei 1 Soma,” and “2 Nuclei 1 Neuron,” provided. **c**. Depictions of classes which are difficult to distinguish, both by human annotations and by the SNIM20 classifier. **d**. Schematic representation of the SNIM20 neural network architecture. Rectangular boxes indicate convolutional layers, where the numbers beneath show the width, height, and number of filters in the respective convolutional layer. Red lines indicate a MaxPool layer. D1 and D2 represent dense layers with 70 and 47 neurons respectively, and an output layer with 20 classes. The table shows an example of the output, where each of the 20 classes is given a prediction score next to its name. In this case, the image was distinctly a single neuron, so SNIM20 assigned a score of 0.987 in the “1 Neuron” class and negligible probabilities in other classes.

### Neuronal Classifier Performance

The training of SNIM20 consisted of 12,689 images across the 20 classes with a train-test ratio of 80:20. We observed that SNIM20 had the capability of distinguishing single neurons (“1 Neuron”) from other classes (AUC=0.99) (**Fig. 3a**). In that same set of images, SNIM20 showed high sensitivity and precision in identifying “1 Neuron” images with a max F1 score of 0.92 (**Fig. 3b**). We assessed the classifier’s performance with its confusion matrix, which further highlighted ability to call the “1 Neuron” class with few errors, sometimes confused with the class “1 Neuron Very Short”, and an overall (multiclass) accuracy of 73.3% for all the classes (**Fig. 3c**). SNIM20 training history shows that by epoch 40, the best weights were found, and the accuracy was maintained across 220 epochs (**Fig. S5**). We next took the prediction score for each of the 20 classes by each of the cells and generated a UMAP representation of the dataset (**Fig. 3d**). Clusters emerged which reflected the true label (although several individual images look out of place). For comparison, and alternative model was also evaluated (**Fig. S6**). We conducted additional validation to assess the accuracy of SNIM20’s predictions in identifying the single neuron class, evaluating SNIM20’s prediction of an independent hand-curated set of 52 distinctly single-neuron images (not part of the original tranche of data). The classifier scored an accuracy of 81% in correctly predicting raft images within the “1 Neuron” class (**Fig. 3e**). 12% (6) of the images were classified as “2 Nuclei 1 Soma” and 8% (4) of the images were unscored.

**Figure 3.**
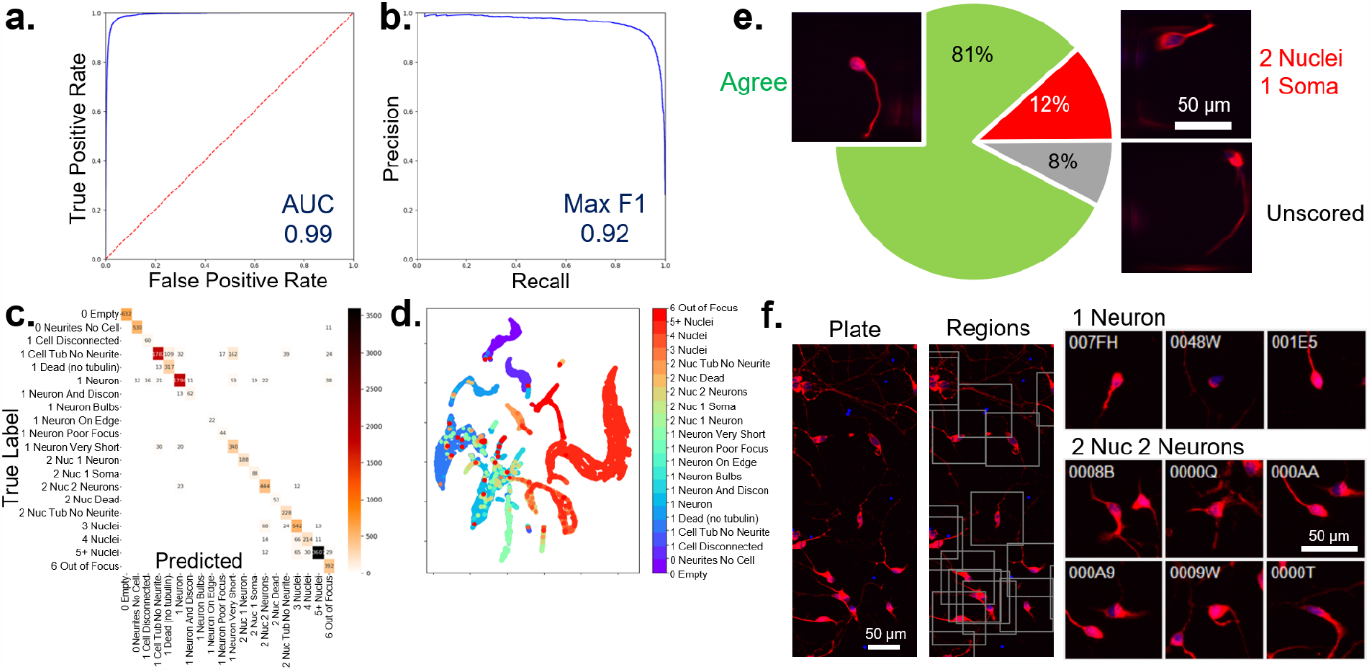
Evaluation metrics and validation of a single-neuron classifier trained from microrafts. **a**. An ROC plot evaluating the “1 Neuron” class versus the other 19 classes for SNIM20. The blue line shows the curve, red is included for reference at the 0.5 level. **b**. Precision-recall curve for SNIM20, “1 Neuron” class (blue line). **c**. Confusion matrix including 20 classes with true labels on the y-axis and predicted labels on the x-axis. The table lists the number of images that fall in each table-cell and is shaded with the heatmap to the right. **d**. UMAP projection of the 20-class prediction scores, colored by the true class label. **e**. Pie chart with the percentage of a new, independently labeled set of images showing those that were correctly labeled (“Agree”), and those that were mislabeled (“2 Nuclei 1 Soma”, and unscored). Example images from the set are included as an inset. **f**. SNIM20 performance on non-microraft plates. Red is MAP2 antibody staining, along with the blue nuclei. Regions defined by the nuclei are outlined by square boxes and are processed like microrafts and automatically annotated by SNIM20. Example predictions of “1 Neuron”, and “2 Nuc 2 Neurons” are shown along with the region IDs in the upper left corner.

The performance of SNIM20 (trained only with neurons on rafts) was further validated in a non-raft setting. On a 96-well plate, SNIM20 identified the neuron or neuron clusters stained for the microtubule-associated marker MAP2 and nuclei (**Fig. 3f**). The fact that the classifier can find many single- and double-neuron ‘regions’ shows that it is flexible to work with different staining and a 96-well plate context, even though it wasn’t trained on either of those. Overall, a CNN classifier (such as SNIM20) can effectively annotate regions for a variety of classes, and has particularly good metrics for identifying single neurons, even in settings that it wasn’t directly trained on.

### Nuclei, Tubulin, and Brightfield Images can each be used to Classify Neurons

Next, we sought to study the relative importance of a set of different wavelengths on the ability to create an effective 20-class model. These are each new models trained on a set of 11,308 images each that contain different wavelengths (**Fig. 4a**), using the pipeline outlined in Fig 1. Instead of just evaluating the best classifier for each wavelength, we examined the trained classifiers to compare their relative performance. Classifiers trained with only one wavelength, either Hoechst (nuclei) or Tubulin (neurites) had the highest validation accuracy, while those trained on Brightfield-only images performed relatively poorly, likely owing to the less specific information provided in that channel (**Fig. 4b**). We also trained classifiers on multi-channel images, so each classifier had access to a combination of two of the channels, or all three (**Fig. 4c**). Classifiers trained on images with both Hoechst and tubulin (as was done with SNIM20) reached the highest overall accuracy, nearing 80%. Interestingly, adding brightfield to the other two channels didn’t increase performance. Also, compared with the single-channel training images in 4b, the multi-channel images had an overall higher accuracy, as expected.

**Figure 4.**
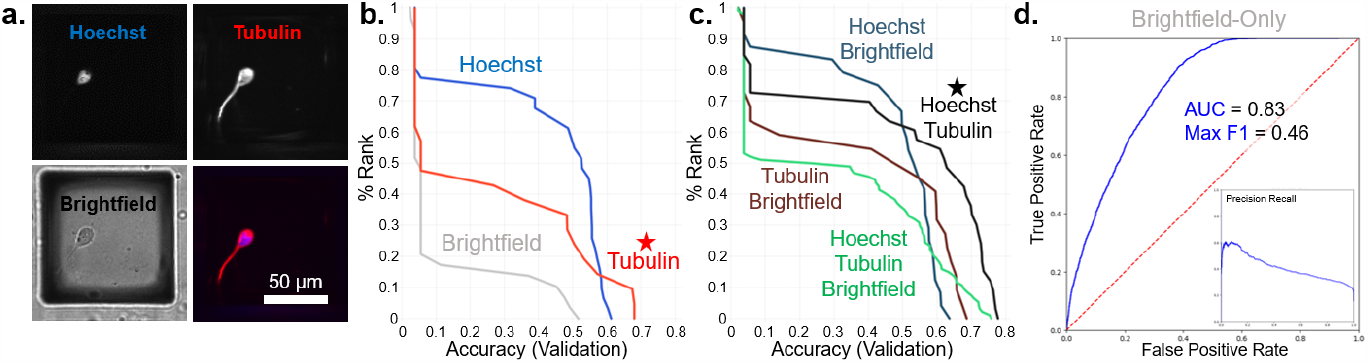
Single and multi-channel training images impact the performance of neuronal classifiers. **a**. Representative images of a single neuron (on a microraft) with two different fluorescent channels, Hoechst for nuclei, tubulin for neurites, and brightfield. Hoechst (blue) and tubulin (red) are shown merged on the bottom right. **b**. Cumulative histogram of multiclass validation accuracies for individual single-channel classifiers, where each curve represents a unique channel: Hoechst (blue), tubulin (red), or brightfield (gray). There are ∽12 classifiers in each curve. The star highlights the tubulin-only classifier as having the highest accuracy. **c**. Cumulative histogram for multi-channel classifiers. The models are grouped by combinations of channels, where the light-blue line represents Hoechst and brightfield classifiers, brown represents tubulin and brightfield, green represents Hoechst tubulin brightfield, and black represents Hoechst and tubulin. There are ∽24 classifiers in each curve. The star marks the curve attaining the highest accuracy. **d**. The ROC (true vs false positive rates) for the highest accuracy Brightfield-only model. The inset shows the precision-recall curve for the same classifier. The AUC and max F1 scores are denoted. The red diagonal line falls at AUC=0.5.

The ability to classify neurons without having to stain or mark the nuclei and neurites is intriguing, so we explored the brightfield-only classifiers in more detail. While the majority failed to successfully classify, the highest-performing brightfield-only model achieved over 50% multiclass accuracy. When comparing between the “1 Neuron” class and other classes, the brightfield-only classifier performed reasonably well (AUC = 0.83, max F1 = 0.46) (**Fig. 4d**). Similarly, another model trained on brightfield-only images also exhibited reasonable discrimination (AUC = 0.78, **Fig. S7**). It is likely that a larger training set could enhance the brightfield-only performance, making it suitable to specific applications where stains or markers are unavailable.

### Coarse and Fine Class Definitions and Model Performance

Being able to distinguish between 20 different classes provides fine detail, but we next asked whether a coarse set of classes changed the classifiers’ abilities. We trained hundreds of individual classifiers by taking the same set of 11,308 images (multi-channel Hoechst and Tubulin) and grouping them into different numbers of classes. For example, in the three-class versions, the “0 Empty” and “1 Neuron” classes were left the same, but the other images were grouped together in a new 3^rd^ class. In the six-class version, the empties and single neurons were again kept un-altered, but other groupings were made. Each classifier always had the exact same number of total training images, but the class definitions were different (**Fig. 5a**). For each individual classifier, we evaluated the AUC of the ROC and the max F1 score comparing the “1 Neuron” class to all other classes, therefore treating the different classifiers equivalently (**Fig. 5b**). We didn’t evaluate the multiclass mean accuracy, since it decreases as the number of classes increases. The high metrics attained with fewer classes is prominent not only in the 3-class versions but also in the 6- and 8-class versions. This is expected since each class now has more training data per class (the total number of images is not divided amongst as many subtly different classes). However, when examining the individual confusion matrices (**Fig. 5c, d**) it becomes apparent that even though the fraction of images confused with “1 Neuron” is lower, the absolute number of confused images is higher. This establishes that various granularities (coarse or fine) of class definitions can lead to reliable image-based classifiers, and that instead, the knowledge provided by the classes should influence the definitions, and not necessarily the overall numbers of classes.

**Figure 5.**
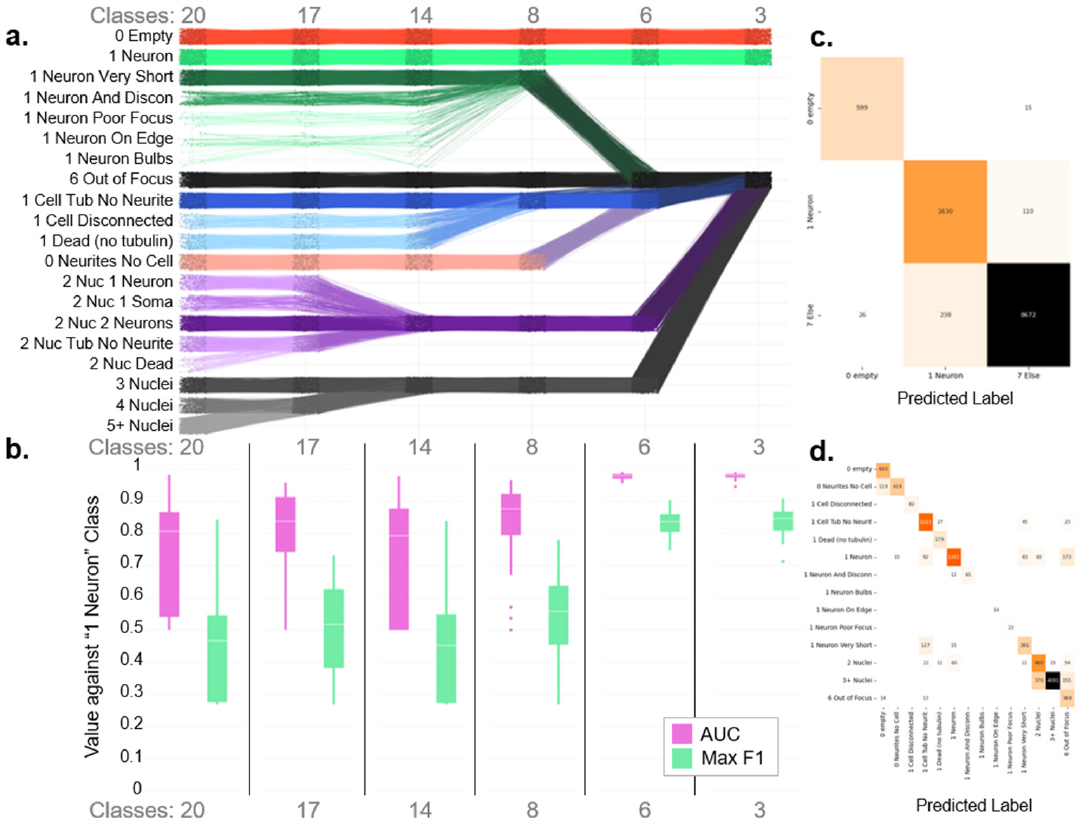
Comparative analysis of classifier performance for differing class number. **a**. A Sankey plot representation of the definitions of the six multi-class granularities from fine (20 classes) to coarse (3 classes). Each horizontal line tracks a single training image (11,308 in total), where its row indicates which grouping is used for that granularity. Convergence points represent the aggregation of images into a single class. Each classifier was trained with the same set of 11,308 images, but these were grouped in different sets of 3, 6, 8, 14, 17, or 20 classes. **b**. Box and whisker plot comparing the AUC and max F1 scores of the classifiers aggregated by class number (granularity) and tested in predicting the “1 Neuron” class. Each column in b. is the same granularity as panel a. above. Pink denotes the AUC score distribution, and green denotes the distribution of the max F1 scores. The white horizontal line within each box represents the median of the distribution, and single points represent outliers. **c**. Confusion matrix of a 3-class version, trained to classify “1 Neuron,” “0 Empty,” or everything else. **d**. Confusion matrix for a 14-class version. Both confusion matrices have true labels on the y-axis and predicted labels on the x-axis. The tables are shaded so darker table-cells indicate higher # of images. Empty boxes represent no recorded instances.

### Custom CNN versus Existing Pre-Trained Transfer Learning Classifiers

Our training pipeline includes the ability to not only randomly sample hyperparameters and neural architectures, but also to replace the initial CNN component of the classifier with existing pretrained models. Therefore, we aimed to optimize the model architecture to assess potential accuracy improvements through transfer learning. We trained hundreds of classifiers, each with the same set of 11,308 images, and each with the original 20 classes from Figure 2. All images contained both Hoechst and tubulin, so the results could be easily compared back to the SNIM20 classifier. The transfer learning models use pretrained weights learned from a different dataset ^16^. We used 7 existing pretrained architectures (ResNet, EfficientNet, MobileNet, NASNet, Inception, Xception, and ConvNexT) as well as our custom CNNs with (AR) and without (NR) residuals. Generally, the use of residuals can improve model performance ^17^. Over two dozen different classifiers were trained with each architecture, and the multiclass validation accuracy varied widely between them (**Fig. 6a, b**). The results showed that the CNN model with residual blocks performed comparably to classifiers that incorporated Xception and NASNet transfer learning layers (medians around 50%, **Fig. 6a**). Some Inception-based classifiers performed very well while others struggled. MobileNet, ResNet, EfficientNet, and ConvNexT had trouble with the fluorescent cellular image data, perhaps because they were trained on mesoscale photographs instead of microscopic images. We also considered the distribution of different classifiers with these architectures (**Fig. 6b**). The custom CNN AR classifier outperformed others, achieving the highest accuracy in this set (SNIM20 would fall in this set). The metrics for that CNN, and some of the best Inception, NASNet, and Xception classifiers show they are capable of high AUC and max F1 scores (**Fig. S8**).

**Figure 6.**
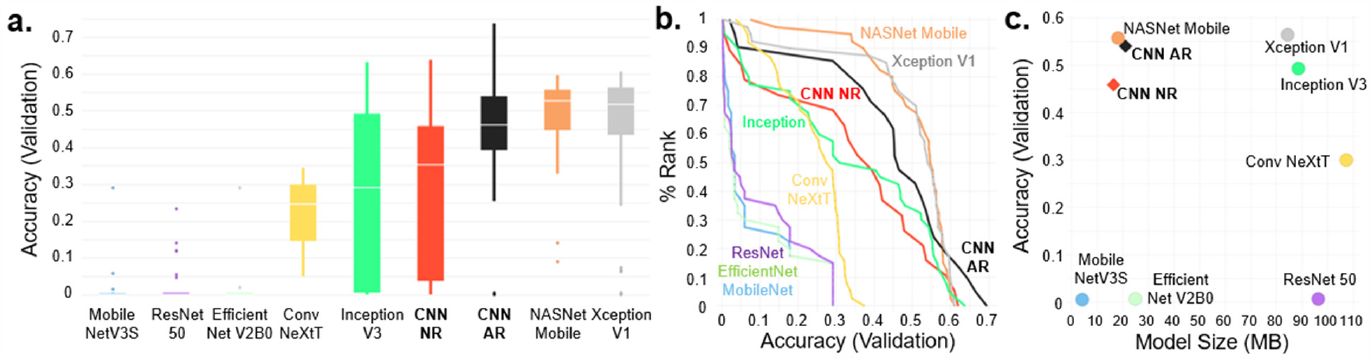
Comparison of custom CNN models with pretrained transfer learning approaches. **a**. Box and whisker plot representing the distribution of multiclass validation accuracy. Each column is a different pre-trained or CNN architecture, with two dozen unique classifiers each. The white horizontal line within each box represents the median of the distribution and single points represent outliers. CNN NR means no residuals were used while CNN AR used residuals. **b**. Cumulative histogram representing the rank distribution of the validation accuracies. Each curve shows a different pretrained or CNN architecture, marked and labeled with distinct colors. **c**. Scatter plot of the average model accuracy as a function of model size in megabytes (MB).

We next wanted to understand the relationship with the model complexity (measured by its size), and their accuracy with the neuronal images. Our CNNs with or without residuals and NASNet Mobile classifiers emerged as having both the highest accuracy while maintaining the smallest size (**Fig. 6c**). While Xception performed well in terms of accuracy, the average size of the trained model was five times larger. A small model size is advantageous due to the less computational costs associated with training and deploying the classifier. Also, the pretrained models here are meant to be ‘generalists’, whereas the custom CNNs are specialists for this particular problem. Hence, starting with NASNet or CNNs with residuals are both appropriate options for these cellular classification tasks.

### Classification Task in Non-Neurons

The training pipeline and model architectures presented are not only useful for neurons, but can also be used more broadly. We therefore sought to develop and test a classifier for a non-neuronal cell type, specifically, human osteosarcoma (U2OS) cells. We followed the pipeline outlined in Figure 1 to identify regions that had single or multiple U2OS cells whose images had nuclear and mitochondrial staining (**Fig. 7a**). In this paradigm, the total number of nuclei in each region (raft) was also separated into classes to distinguish high quality single-cell rafts “1 Cell”, from other cases, using 12 classes total (**Fig. 7b**). After the refinement round with just 2,007 training images, we chose a “SCIM12” classifier that had high accuracy (89.1%) and very little confusion between classes (**Fig. 7c**). When examining the details of how well this model performed in differentiating “1 Cell” with all the other classes, the metrics were very high (AUC = 0.99, Max F1 = 0.95, **Fig. S9**). As in Figure 4, we also trained classifiers that had only Hoechst or only mitochondrial channels and compared them to the images with both channels (**Fig. 7d**). The models with both Hoechst and mitochondria performed the best, with accuracies approaching 90%. Therefore, the pipeline presented in this manuscript can give high quality annotations for both neuronal and non-neuronal contexts.

**Figure 7.**
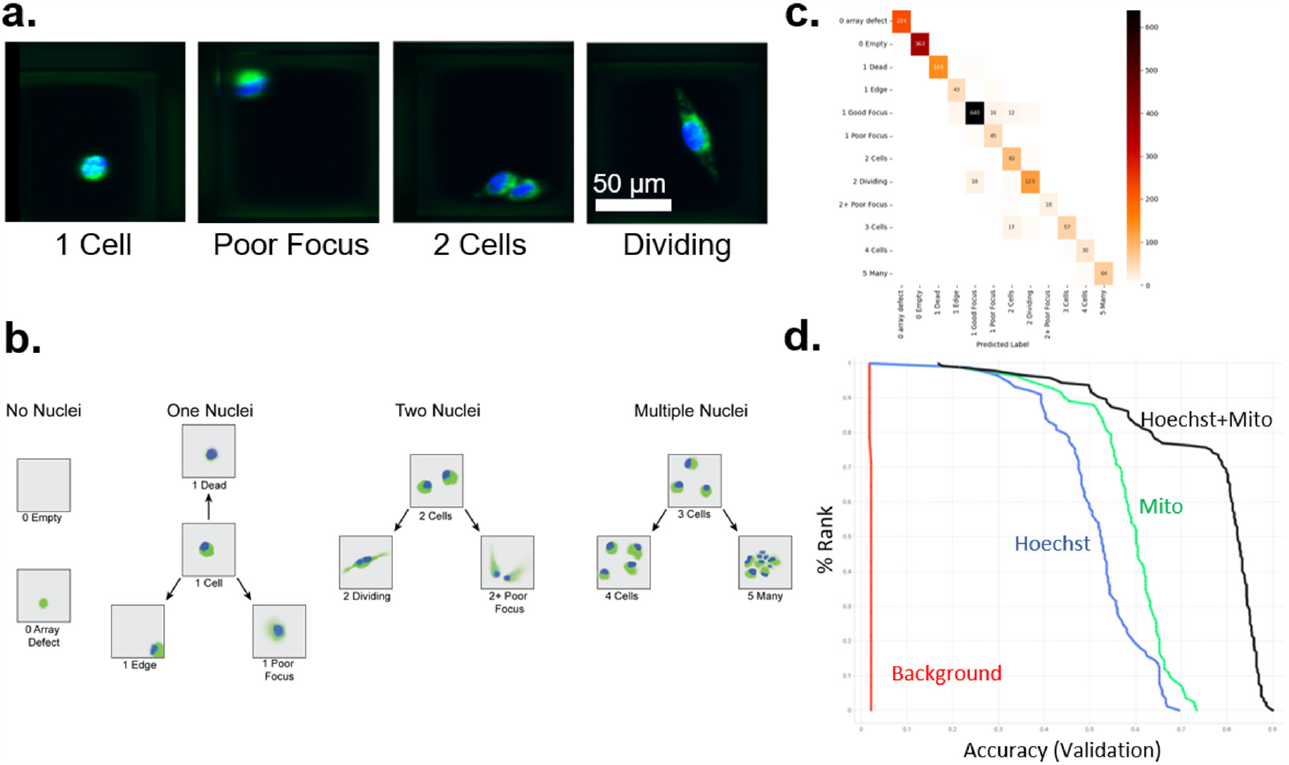
Automatic annotation of healthy U2OS cells. **a**. Representative confocal microscope images for U2OS cells stained for nuclei (Hoechst in blue), and mitochondria (green). Examples of the “1 Cell,” “Poor Focus,” “2 Cells,” and “Dividing” classes are shown. **b**. The U2OS images were categorized into four groupings, those with no nuclei, one nucleus, two nuclei, and multiple nuclei. The 12 specific classes are shown in cartoon form. **c**. Confusion matrix for the SCIM12 classifier, with true labels on the y-axis and predicted labels on the x-axis. The numbers in each table-cell indicate the number of images and are shaded according to the heatmap on the right. **d**. Cumulative histogram for a series of trained classifiers solely with Hoechst, mitochondria, or both are shown as blue, green, or black curves. The multiclass validation accuracy is shown on the y-axis. The red line indicates a background set for comparison.

## Discussion

*Neuronal* classification and segmentation tasks pose numerous challenges owing to inherent complexities of neuronal cultures, characterized by their tendency to cluster, thrive in high density, and exhibit complex morphologies. In massively parallel approaches which require single cell capture and phenotypic analyses, such factors present significant hinderances for identification and capture of single neurons. Furthermore, existing algorithmic frameworks fail in such tasks, partially due to high variability across neuronal images. However, deep learning tools are well suited for such tasks, which we demonstrate in “SNIM20”, a CNN classifier for neuronal image annotation. Since RaftSeq and other morphology-based functional genomics platforms require a rapid categorization of a large volume of images to identify single neurons, developing classifiers such as SNIM20 and its variants can address these concerns. As an open-source and versatile tool, this framework presents a more accessible and customizable tool for semi-automated image classification ^18,19^.

We opted to use a CNN framework because it presents a good tradeoff between quality and training size and is commonly used in image recognition, segmentation, detection, and information extraction ^20^. To enhance the model’s performance, we varied the CNN architecture and the number of fully connected dense layers. Additionally, we fine-tuned the model by randomizing hyperparameters, only selecting the best classifier from a series. These optimizations allowed us to fine-tune the classifier while also mitigating overfitting, leading to favorable results for the identification of single neurons. SNIM20 is an example of a classifier that can annotate differentiated neurons and rapidly determine which are the optimal ones for downstream study. Other researchers can apply this pipeline to identify single neurons or other cell types from microscopic images, streamlining the analytical process. Despite the training set for these classifiers being from microraft cultures, they can be trained and inferred on more common plate-based formats (**Fig. S2**).

We have recently started initiatives for two new applications of the classification system. We plan to hand classify co-cultures of microglia and neurons. In this context, our objective is to categorize microglia based on their ‘neighborhood status’, discerning whether microglia are solitary or in proximity to neurons. In another application, we plan to examine adipocytes on a feeder layer of mesenchymal cells, where we will hand-annotate adipocytes based on the intensity of their respective lipid droplets. In both cases, a single expert annotator can then be virtually ‘copied’, so that the classifier can automatically give similar classifications in a manner only biased by the original training set, but not by different human labelers, as would otherwise be necessary.

Another important future aim is to address neurite disentanglement directly. While the raft-based screening keeps the neurites separated, most other contexts include highly overlapping and complex neurite arbors. The development of a disentanglement-focused model will enable users to analyze images featuring multiple neurons, even without ‘rainbow’ stains like BrainBow or TetBow ^21^. This approach would effectively address the limitations inherent in many current methodologies ^22–25^. With the expansive potential of a CNN-based classification system, we can harness is to advance the field of biomedical image analysis and examine individual cells in detail in a high throughput manner.

## Methods

### Human iPSCs Maintenance and Neuronal Differentiation

The neuronal cells were derived from the WTC-11 human induced pluripotent stem cell (iPSC), some with the addition of an inactive gRNA library for downstream experiments. The iPSCs were obtained from Genome Engineering and Stem Cell Center at Washington University in St. Louis, propagated in mTeSR plus media (100-0276, StemCell Technologies, Vancouver, Canada), and seeded on 6-well plates coated with Matrigel (354277, Corning, Corning, USA), diluted 1:6 in DMEM/F12 media (11320033, Thermo Fisher, Waltham, USA). The iPSCs were maintained, expanded, and incubated at 37°C and 5% CO_2_. The media was replaced every day and cells were passaged once they reached 80% confluency.

Cells were rinsed with Dulbecco’s Phosphate-Buffered Saline (DPBS, 14190144, Thermo Fisher) and treated with 1mL ReLeSR (05872, StemCell Technologies) at room temperature for one minute. ReLeSR was then aspirated, and cells were incubated in a 37°C incubator for 5 minutes before being collected and split at a 1:10 dilution in a new plate. Upon reaching 80% confluency, iPSCs were differentiated into motor neurons as per the protocol outlined by Du et al ^11^. In brief, the natural developmental signaling factors were recapitulated with small molecules including Ascorbic acid (72132, StemCell technologies), CHIR99021 (72054, StemCell Technologies), DMH-1 (73634, StemCell Technologies), SB-431542 (72234, StemCell Technologies), Retinoic acid (R2625, Sigma, St. Louis, USA), Purmorphamine (72204, StemCell Technologies), ROCK inhibitor (Y-27632, 72304, StemCell technologies), and Compound E (73952, StemCell Technologies) using the protocol ^26^. Additionally, the differentiation media also contained N-2 (17502001, Thermo Fisher) and B-27 (A3582801, Thermo Fisher) supplements. iPSCs underwent differentiation into motor neurons across four distinct stages, incorporating varying time and concentrations of the various small molecules (**Fig. S1**).

The induction of neuroepithelial cells (step 1) was initiated by introducing stage 1 differentiation media composed of 50% neural basal (21103049, Thermo Fisher), 50% DMEM/F12, 0.5X N2, 0.5XxB2, 0.1mM Ascorbic acid, 1X Glutamax (35050061, Thermo Fisher), 1X 0.1% Penicillin/Streptomycin (15140122, Thermo Fisher), 2 μM DMH-1, 2 μM SB-431542, and 3 μM CHIR99021. On day 7, the cells were then transitioned to stage 2 media, introducing two additional small molecules, namely, 0.5 μM Purmorphamine and 0.1 μM Retinoic acid, while reducing CHIR99021’s concentration from 3 μM to 1 μM. This stage induces OLIG2+ expression, initiating the cells’ motor neuron progenitor (MNP) identity. On day 4 of being on stage 2 media, about 1 million cells were frozen down per aliquot for later experiments (see below, raft plating and imaging).

Between each stage, cells were dissociated with TrypLE (354277, Gibco, Gaithersburg, MD, USA) and re-plated. Each time the cells underwent the differentiation process and required dissociation; they were treated with 10mM ROCK inhibitor (72304, StemCell Technologies) for 24 hours following the passage. For dissociation, cells were washed with DPBS, 1X TrypLE was introduced, and the vessel was incubated at 37°C for 3-5 minutes. The cells were then centrifuged at 200*g* for 5 minutes before resuspending in appropriate media. Noteworthily, while resuspending the cell pellet, several triturations were performed to break the cell clusters into single cells.

### Raft Plating and Imaging Motor Neurons

MNPs were seeded on two different kinds of raft plates, a single reservoir and a quad reservoir, each containing 100×100μm microrafts (Cell MicroSystems, Inc., NC, USA). To prepare the plates for cell seeding, the plates were washed with 1 mL/well and 5 mL/well (for quad and single raft plates, respectively) of 1xPBS (10010023, Corning) and incubated at 37°C for 5 minutes. These washes were done three times to remove the glucose layer from the raft plates. The raft plates were then coated with 4μg /mL Poly-d-lysine (PDL, P0899, Sigma) and incubated overnight at 37°C. The next day, the raft plates were rinsed three times with molecular grade water (46-000-CV, Corning) and coated with 5μg/mL mouse laminin (23017015, Fisher Scientific) for similar overnight incubation. Both PDL and Laminin solutions were made with molecular grade water.

MNPs frozen on stage 2 day 4 were thawed, centrifuged at 200*g for five minutes*, resuspended in 1mL stage 2 media, and subsequently plated on a Matrigel-coated 6-well plate. The cells are then grown for an additional 3 days on stage 2 media. On the 14^th^ day since the start of the differentiation process, the cells could be nucleofected with Cas9 plasmid and plated on stage 4 media consisting of 50% neural basal media, 50% DMEM/F12, 0.5X N2, 0.5X B27, 0.1mM Ascorbic acid, 1X Glutamax, 1X 0.1% Penicillin/Streptomycin, 0.1 μM Purmorphamine, and 0.5 μM Retinoic acid. Approximately 2 million cells were nucleofected in a large cuvette per reaction using Lonza’s P3 primary cell 4D-nucleofector kit (V4XP-3024, Lonza, Basel, Switzerland). The cells were allowed to recover from the transfection for 3 days while inducing MNX+ motor neuronal cell fate. In this manuscript, Cas9 plasmid was not added, but this detail is included for more clarity on the overall timing.

On day 18 of differentiation, the cells are seeded on raft plates. For quad and single raft plates, 80,000 cells/well and 320,000 cells/well are seeded respectively. Cell counts for raft plating are performed using a hemocytometer (with higher accuracy than automated cell counters). The next day, media is changed to stage 5 media, which consists of stage 4 media with the addition of 0.1 μM Compound E. Notably, the protocol mentions stage 3, an optional MNP expansion step, which we skip. Full media changes are conducted every day during stages 1, 2, and 4, and a half media change every other day when on stage 5.

Motor neurons are stained and imaged 48-hours after the introduction of stage 5 media. A master mix of the staining media is prepared, consisting of 4.05 μM Hoechst 33342 (H3570, Thermo Fisher), and 1 μM Tubulin Tracker Deep Red (T34076, Thermo Fisher). For staining, the media is gently removed from the plates, and 500μL/well and 2mL/well of stage 5 media is added to the quad and single plates, respectively. Following the application of the stain, the plates are incubated at 37°C for 30 minutes, after which the stain media is removed and three washes with Step 5 media in the same volume as the stain media are performed. For imaging, the plates are positioned on a Cell Microsystems plate adapter and imaged on Molecular Device’s INCell 6500HS Confocal microscope at 20x with 0.45 NA. The live cell chamber is used, set to 5% CO_2_ and 37°C. The imaging protocol included 40,000 rafts / 1,368 fields and 25,600 rafts / 960 fields in the single and quad plates, respectively, with 12% field of view overlap. Exposure times for Hoechst (405 nm), Tubulin Tracker Deep Red (642 nm), and brightfield all averaged less than 100ms.

### Immunofluorescence to Confirm Motor Neuron Identity

To determine the iPSC’s differentiation proficiency, cells are analyzed with immunofluorescent staining at different stages during the differentiation process. First, cells are fixed with 4% paraformaldehyde (PFA, 158127, Sigma Aldrich) for 15 minutes and washed three times with 1X PBS. Next, cells are permeabilized using 0.5% Triton-X 100 (93443, Sigma Aldrich) for 15 minutes and blocked with 3% Bovine Serum Albumin (BSA, A9418-10, Sigma-Aldrich,) for 1 hour. Both primary and secondary antibodies are diluted with BSA, placed on cells and incubated for 1 hour. The primary antibodies tested are 1:75 SOX1 (AF3369, R&D Systems), 1:500 Olig2 rIgG (AB9610, Sigma), 1:100 NKX2.2 mIgG (74.5A5, DSHB), 1:75 MNX1 mIgG (81.5C10, DSHB), 1:70 ISL1 mIgG (40.2D6, DSHB), 1:300 CHAT gIgG (AB144P, Sigma-Aldrich), and 1:100 MAP2 rIgG (MAB3418-25UG, Thermo Scientific). The secondary antibodies were used: 1:1000 donkey anti-mouse (A-31571, Thermo Fisher), 1:1000 donkey anti-rabbit (ab150075, Thermo Fisher), and 1:200 donkey anti-goat (A-21432, Thermo Fisher). Through confocal microscopy (INCell Analyzer 6500), the cells are then imaged, and cell tracing and fluorescence intensity extraction was done using IN Carta Image Analysis Software (Molecular Devices).

### Passaging and Maintenance of U20S Cells

In some experiments we utilized a human osteosarcoma cancer cell line, U2OS cells (HTB-96, ATCC). The U2OS cell line engineered to express doxycycline-inducible Cas9 was acquired from GESC@MGI WashU. The U2OS cells were cultured in McCoy’s 5A Modified Medium (16600082, Gibco) supplemented with 10% tetracycline-free fetal bovine serum (FBS, PS-FB3, Peak Bio, Pleasanton, CA, USA). Cells were grown in either T75 or T150 tissue culture flasks and were passaged at 90% confluency. Trypsinization consisted of placing 0.25% Trypsin-EDTA 1x (25200056, Gibco) on the cells for 5 minutes at 37°C, followed by collection of cell suspensions and centrifugation for 3 minutes at 1200 RPM.

### U2OS Cell Imaging

The U2OS cells were plated on raft plates, either a single plate divided into 40,000 microrafts or a quad plate divided into 25,600 microrafts (Cell Microsystems). 48,000 U2OS cells were seeded in a single raft plate, while 12,000 U2OS cells were seeded per well in the quad plates. Cells were plated 48 hours prior to staining with 4.05 μM Hoechst, and 0.5 μM MitoTracker Green (M7514, Thermo Fisher) for 30 minutes at 37°C. Like neuronal imaging, U2OS imaging consisted of placing the plates in a Cell MicroSystems plate adapter and imaging on the Molecular Devices INCell 6500HS Confocal microscope at 20x with 0.45 NA. The live cell chamber was utilized, set to 5% CO_2_ and 37°C. In total, 40,000 and 25,600 fields were imaged in the single and quad plates respectively, with 12% field of view overlap. Exposure times for Hoechst (405 nm) and MitoTracker Green (488 nm) averaged < 100ms.

### Training Image Annotation and Sorting

Some of the general details pertaining to image annotation via FIVTools are listed in and illustrated in **Figure S2** ^1^. The 16-bit multichannel images of single 20x fields-of-view are loaded directly into FIVTools. If the cells are plated on microrafts, then raft calibration is performed (which connects the plate coordinates to raft coordinates). For 96-well plate experiments, nuclei segmentation is performed first, to identify the raft-sized regions around each nucleus. Image-display parameters are then adjusted to optimize the contrast for both nuclear and accessory channels. The neuronal classifiers were trained with nuclei in the blue channel and tubulin in the red channel. U2OS classifiers were trained with nuclei in the blue channel and mitochondria in the green channel. It is worth noting that these channels can be swapped as needed. Initially, half a dozen experienced scientists manually annotated each of the scanned raft images in a well by pressing a number key (which designates the class index, initially arbitrary), then clicking on a raft to assign that class label (see **Fig 2** for the list of labels). These labels can be checked using the “Check Annotation” feature in the program, enabling the user to quickly visualize groups of images from different classes. This functionality provides flexibility, allowing users to modify annotations at any time. Next, all the annotated regions are exported with the adjusted parameters, so that the images are placed in a sub-folder with the name of the class. These images are exported as RGB 8-bit per channel bitmaps with a width and height of 128×128 pixels. The number of images per class are listed in **Table S1**.

### Machine Learning Training

We used Keras/Tensorflow 2.10 to train the CNNs. The workflow was executed in a Jupyter notebook inside of Visual Studio Code, using a python kernel in Anaconda. See “TF_NB_MainCells01.ipynb”, then navigate to the code section “Raft CNN – Classifiers” > “Directory Defines Classes” > “RGB 8-bit per Channel”. First, images were loaded into a TF dataset, cached, and prefetched. Each classifier was generated using “neural architecture search”, by randomizing various parameters and then inferring on additional images. The following parameters were randomly assigned: CNN kernel size (2-4), CNN initial number of filters (18-36), CNN depth (max(1, log(image width, kernel size))), and filter increase rate (1.25 to 1.75). The number of dense layers after convolution starts with a random size that is no more than half of the flattened size. In addition, the dense layer dropout is randomized, and whether or not to do batch normalization in the dense layers, residuals in the CNN layers, and whether to use single loss or multi loss for each class is randomly turned on and off.

Images were resized, rescaled between 0 and 1, then fed through the convolutional layers or the transfer learning layers (more below). Early stopping was used if the loss stopped decreasing, usually with a patience set at ten percent of the number of epochs, which was usually set to be around 200. Batch size was 64, and training:test split was 80:20. As soon as each training finished, the script exported several evaluation metrics, including the best loss and accuracy as well as inference on the full dataset. Then the thread was reset, a new random set of parameters was created, and the process was repeated. Over a hundred individual classifiers were automatically and independently trained in each round.

Most of the manuscript did not use transfer learning with pre-trained layers, but this was implemented for one specific experiment (**Fig. 6**). In that case, a variety of transfer layers used included Keras’s built in convnext.ConvNeXtTiny, efficientnet_v2.EfficientNetV2B0, inception_v3.InceptionV3, MobileNetV3Small, nasnet.NASNetMobile, ResNet50, and xception.Xception instead of the CNNs mentioned above.

### Evaluation and Analysis

In addition to assessing the loss and the accuracy of each classifier, we also treated the entire full inferred output as two classes to calculate additional metrics. Given that both the classifiers, neuronal and U2OS, had one ‘ideal’ class that was the purpose of the classifications, we reduced the labels to represent just the preferred class (1 Neuron or 1 Cell) vs all other classes. Afterwards, we computed the AUC for sensitivity/specificity and the Max F1 score. Figures and results were visually represented using either the software Spotfire (Tibco, California) or through Matplotlib in Python.

## Addenda

### Author Contributions

P.P., L.K.MA., M.G.W., K.P.G., J.E.W., C.L.K., and W.J.B. wrote the manuscript. W.J.B., P.P., J.E.W., and C.L.K. designed the experiments. Initial planning and ideas brought by W.J.B., P.P., J.E.W., and C.L.K.

P.P., C.L.K., J.E.W., L.K.MA., M.G.W., K.P.G., U.K., S.N.E. performed the experiments. U.K., L.K.MA., W.J.B., designed the figures. P.P., L.K.MA., G.W.B., and W.J.B. analyzed the results. W.J.B. wrote some of the key software. All reviewed the paper and gave suggestions.

## Acknowledgements

We would like to thank Jack Bramley, who started early versions of this in 2019. Thanks to Diana Grigore and Josh Milbrandt for their help with aspects of the project. We would also like to thank Jeffrey Milbrandt and Robi Mitra for their continued support. We would like to thank the **McDonnell Genome Institute**, specifically GESC@MGI. Finally, we would like to thank the Genetics and MGI administration and our maintenance and cleaning staff.

## Code Availability

https://gitlab.com/buchserlab/fivetools

https://gitlab.com/buchserlab/five_notebooks

