## Supplemental Figures for "Classification of iPSC-Derived Cultures Using Convolutional Neural Networks to Identify Single Differentiated Neurons for Isolation or Measurement"

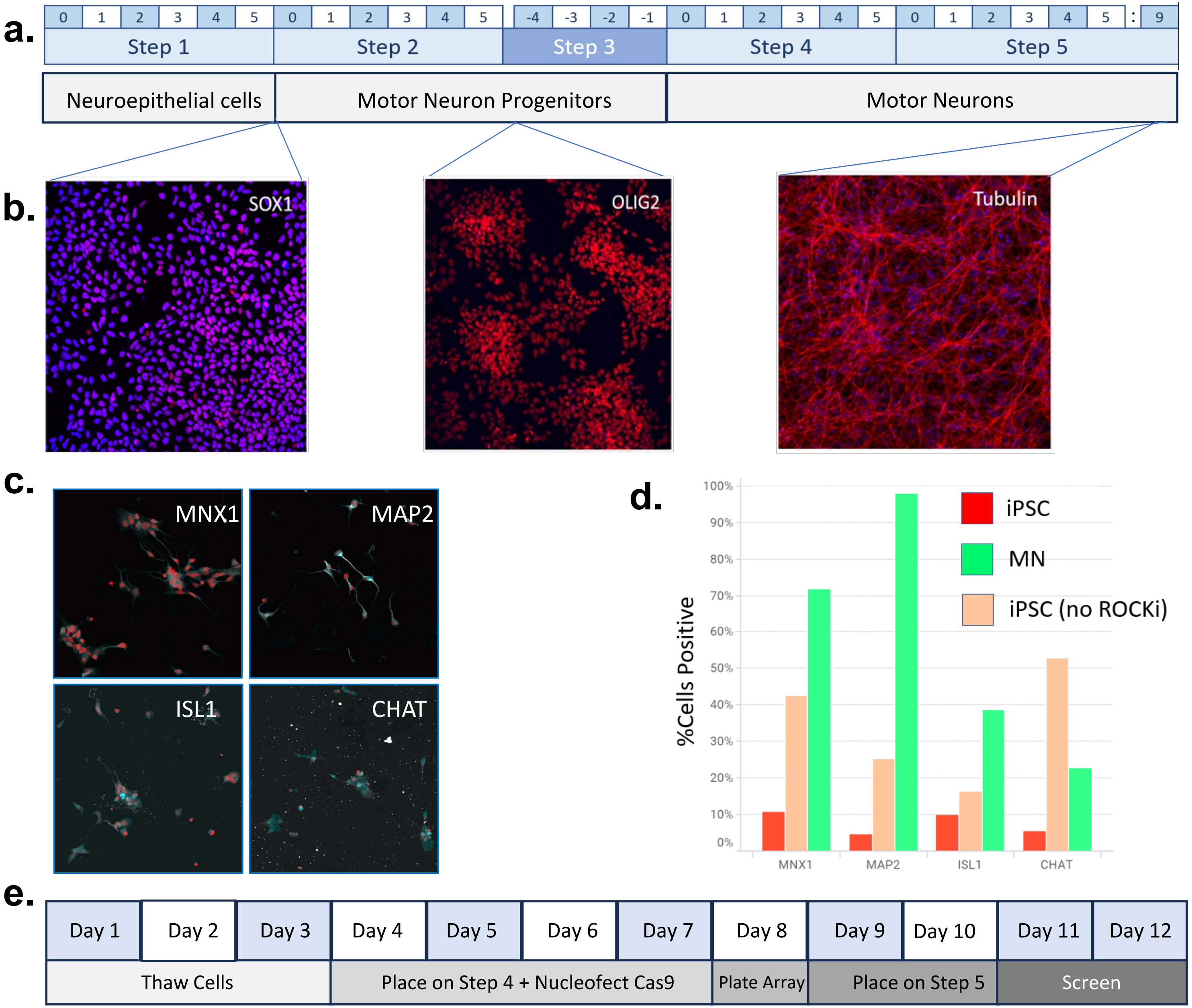


**Figure S1. iPSC-to-motor neuron differentiation and screening pipeline. a.** Table showing the number of days within each step, as well as the respective cell type. **b.** Immunofluorescence staining shows markers with expected expression at the respective steps. **c.** Immunofluorescence staining was done at Step 5 to confirm the expression of MNX1, MAP2, ISL1, and CHAT, all characteristic of mature motor neurons. **d.** Bar charts quantifing the percent of cells positive for each stain in c. **e**. Schematic with the optimized screening schedule beginning with Day 1, where Step 2 Day 4 cells are thawed and culminating on Day 12, the second day of the screen.

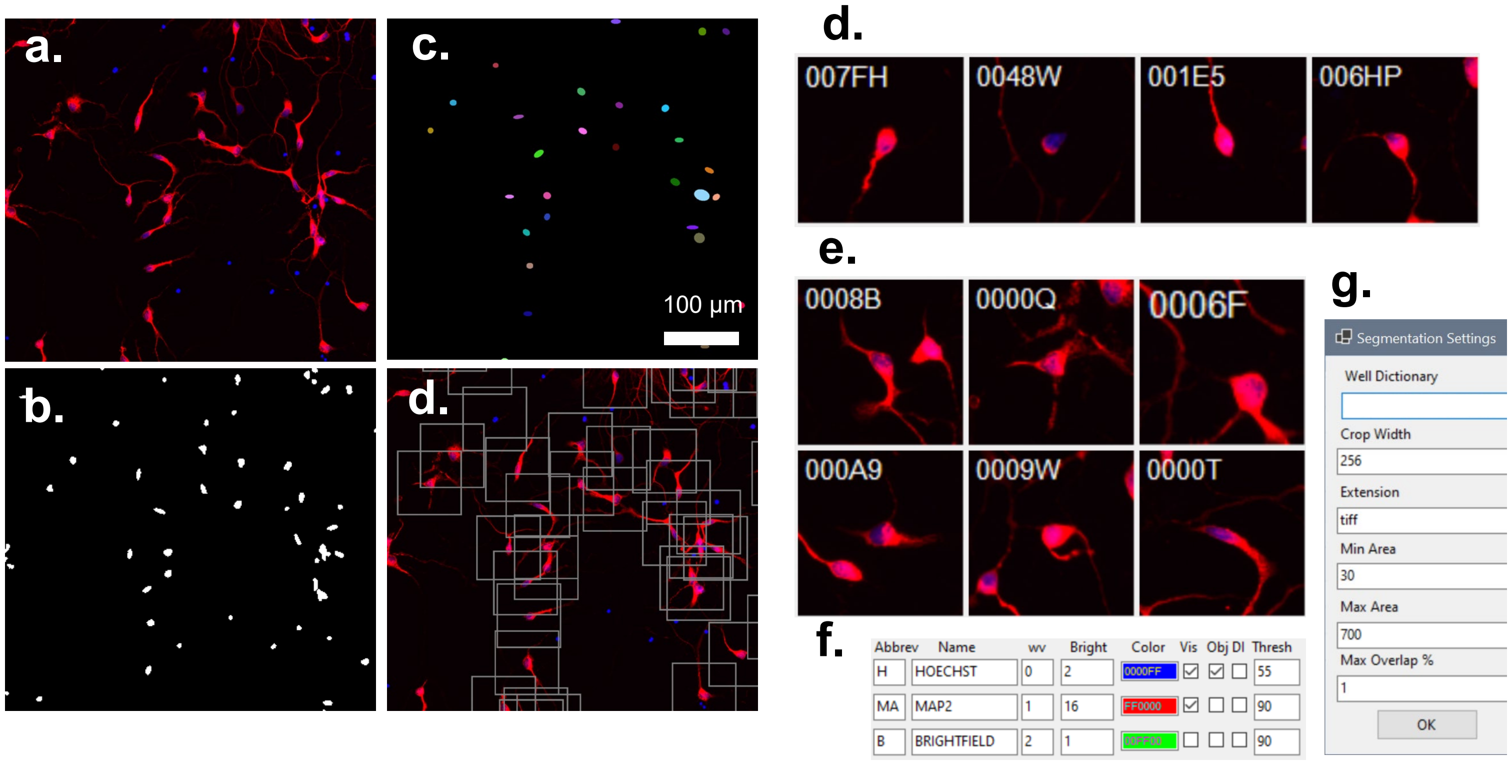


**Figure S2. Plate-based regional analysis overview.** First, use FIVTools Image Check to adjust wavelength intensities to your liking. For neuronal classification, the nucleus should be in the blue channel and the neurites should be in the red channel (**a**). Set a threshold between 1 and 255 (90 is default), then double click the threshold to preview the threshold (**b**), object identification (**c**), and regions (**d**). Then, the label "classification" can be right clicked to choose a classification model, and the left clicked to run classifications. After classifications, FIVTools "Check Annotations" can be run which shows the various classified regions, in this case single neurons (**d**), and double neurons (**e**). Various settings are adjustable, including the wavelengths, intensities, colors, and thresholds (**f**), and the region size (aka Crop Width), the object min and max area, and the % overlap (where 1 means take all, and 0 means disallow overlap) in panel **g**.

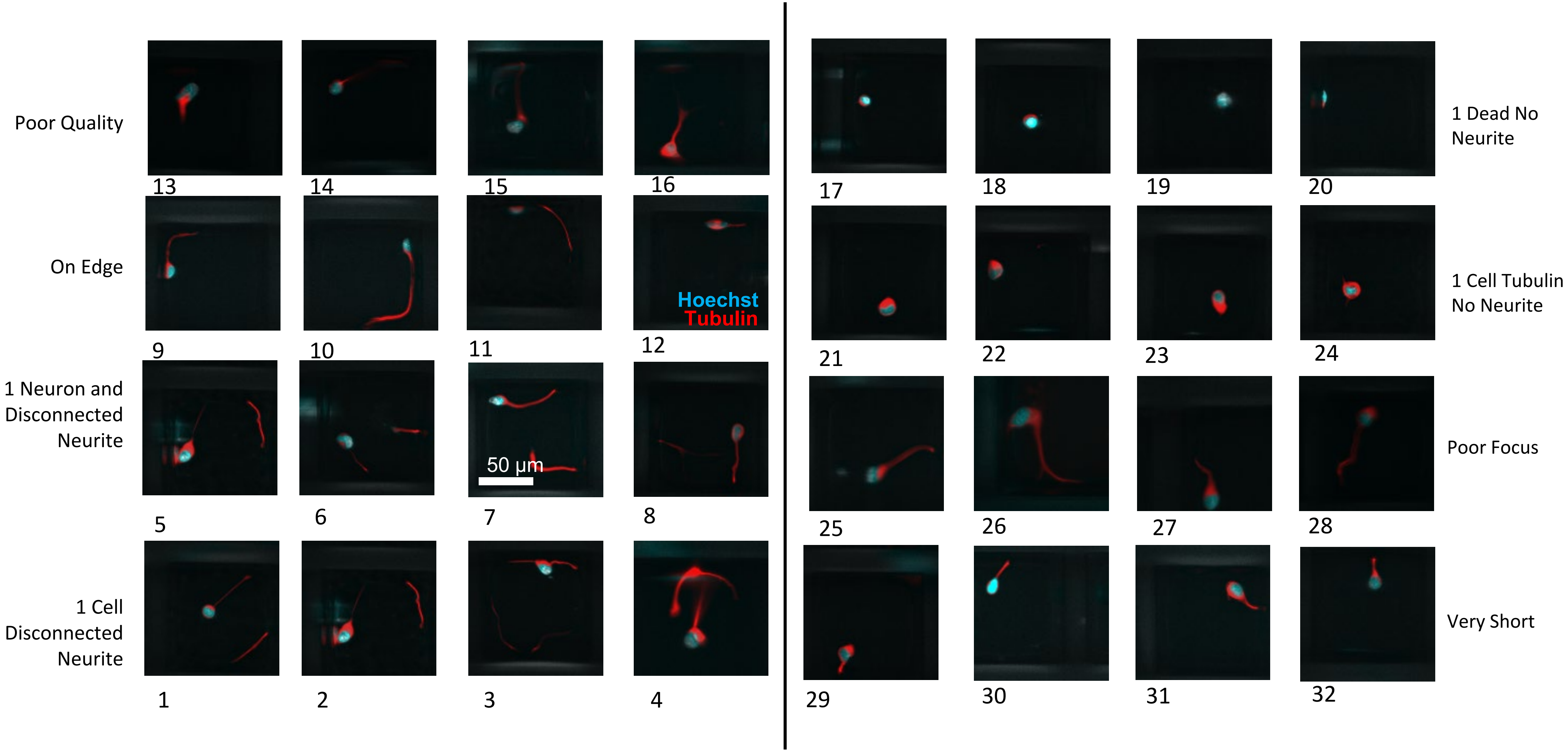


**Figure S3. 1 Neuron Additional Classes.** Additional 20x confocal microscopy images of the classes represented under the "1 Nuclei" grouping. Red is Tubulin, Blue is Hoechst, and white is Hoechst over Tubulin. Names near each row indicate the class of the respective images.

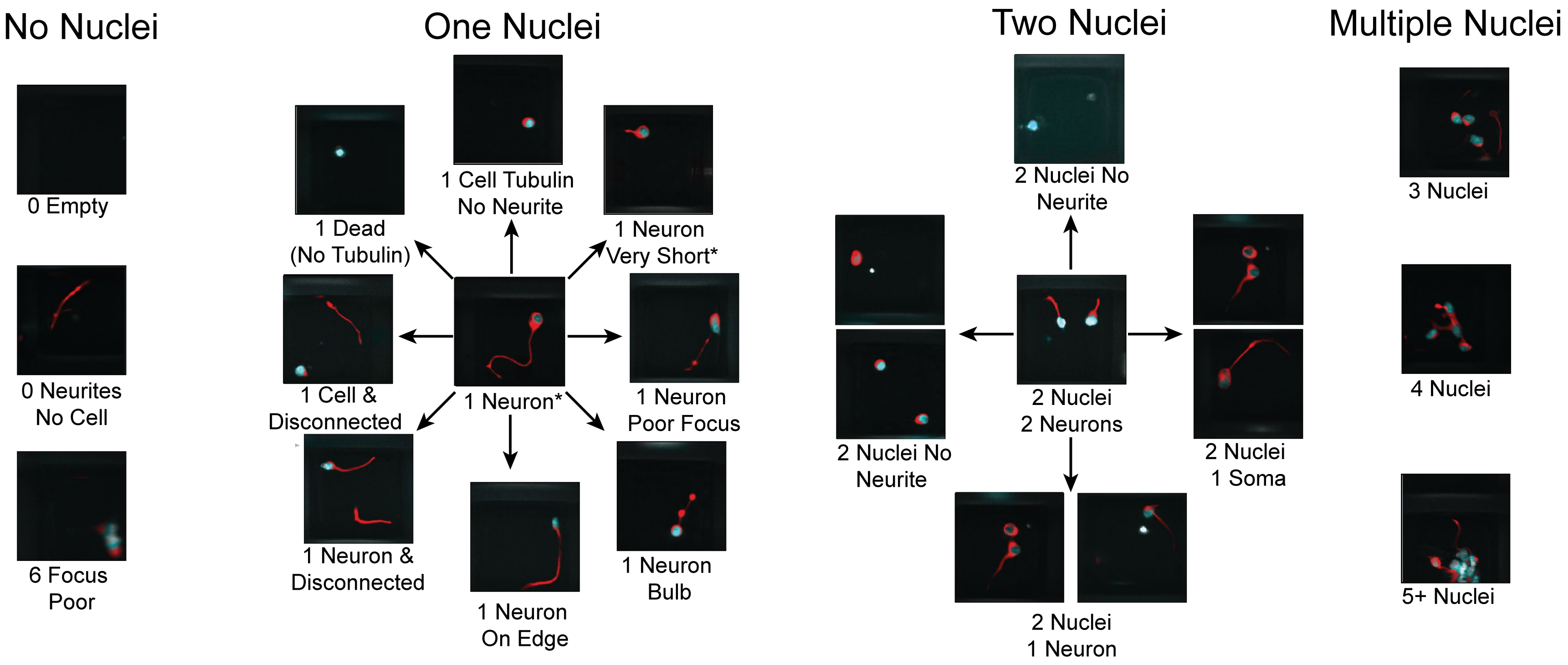


**Figure S4. Physical representation of classes.** 20x confocal microscopy images of each of the 20 image classes under the four groupings, No Nuclei, One Nuclei, Two Nuclei, and Multiple Nuclei. Each square represents a microraft. Red is tubulin, blue is Hoechst, and White is Hoechst over Tubulin. Names under the microrafts identify the class which the respective image represents. Refer to Figure 2.

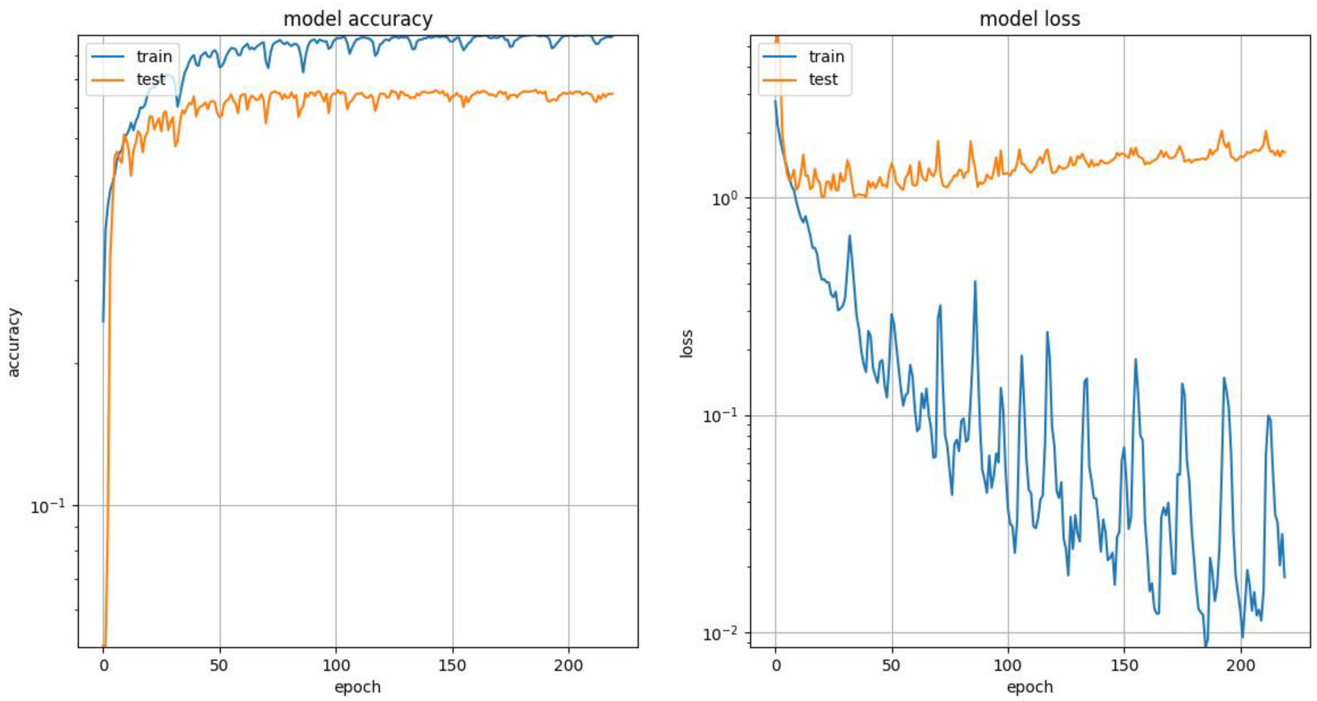


**Figure S5. Training history for SNIM20.** On the left shows the accuracy per epoch of the training (blue), and testing (orange) datasets. On the right shows the loss for the same sets. Accuracy and loss both plateau around epoch 20, while some the ideal set of weights is found at 40 epochs. Full name of the model is acc_0.7371 CNN_Ar CNNs_8 kn_2 flt_33 iRt_1.25 ds__70, 47_ nE_220,2200 bSz_64 bNm_1 i_23.

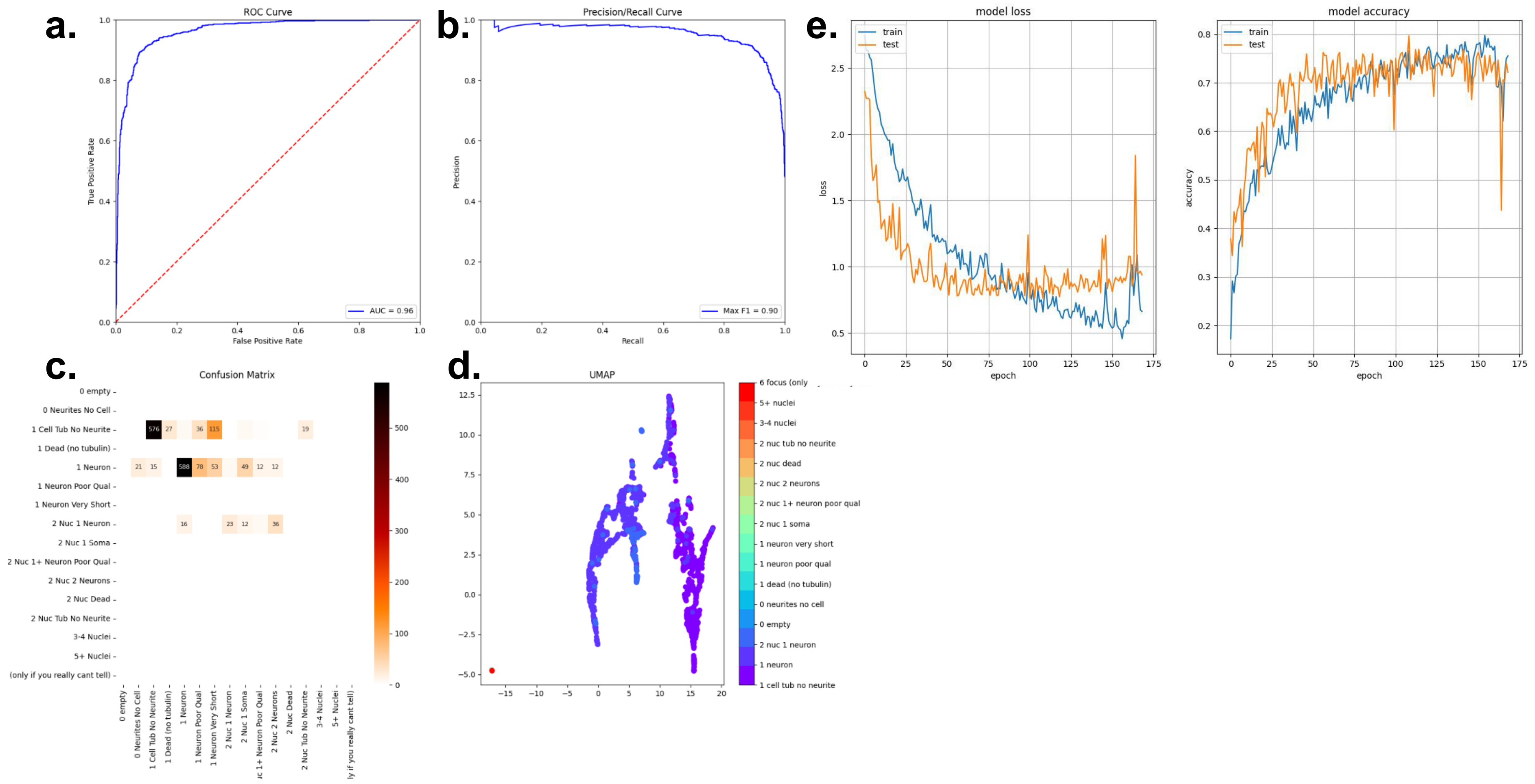


**Figure S6. SNIM16 Neuronal Classification.** SNIM16's accuracy in predicting the "1 Neuron" class versus "1 Cell Tub no Neurite" and "2 Nuc + 1 Neuron" a. AUC ROC curve. B. F1 score (precision/recall) curve. C. Confusion matrix and D. UMAP representation for SNIM16's class predictions. E. Model loss and accuracy curves, where blue represents the train set and blue represents the test set.

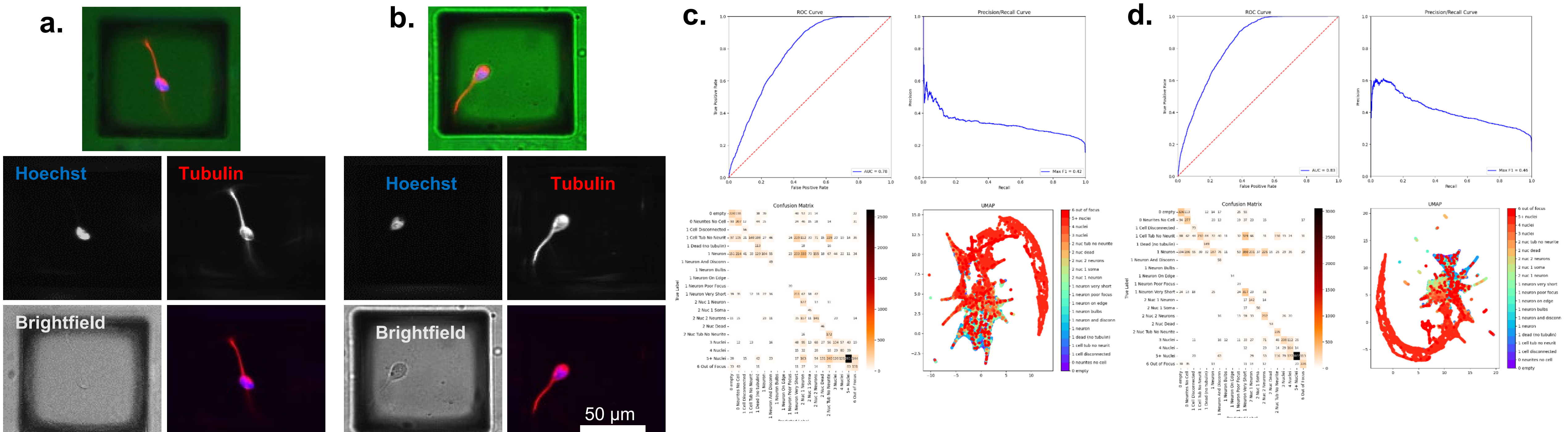


FIV831A2_4.A - 2.179.N8P6.0

FIV773A2_1.B - 2.55.J9F3.0

ev acc_0.3437 c1 CNN_Ar CNNs_6 kn_2 flt_32 iRt_1.75
ds__1000, 518, 74, 69_ nE_160,1600 bSz_64 bNm_0 i_26

ev acc_0.4159 c1 CNN_Ar CNNs_7 kn_2 flt_24 iRt_1.5
ds__70_ nE_160,1600 bSz_64 bNm_1 i_15.png

**Figure S7. Fluorescent Channels and Best Brightfield-Only Classifiers.** a, b) Example micrographs from two distinct rafts with single neurons on them, showing the nuclei in Hoechst, the tubulin with Tubulin tracker Deep Red, and the brightfield channel (raft boundaries are easily discernible). c, d) the two best brightfield-only classifiers (named below the figure). Each shows the ROC, Precision/Recall curves on the top, and a confusion matrix and a UMAP for all 20 classes on the bottom.

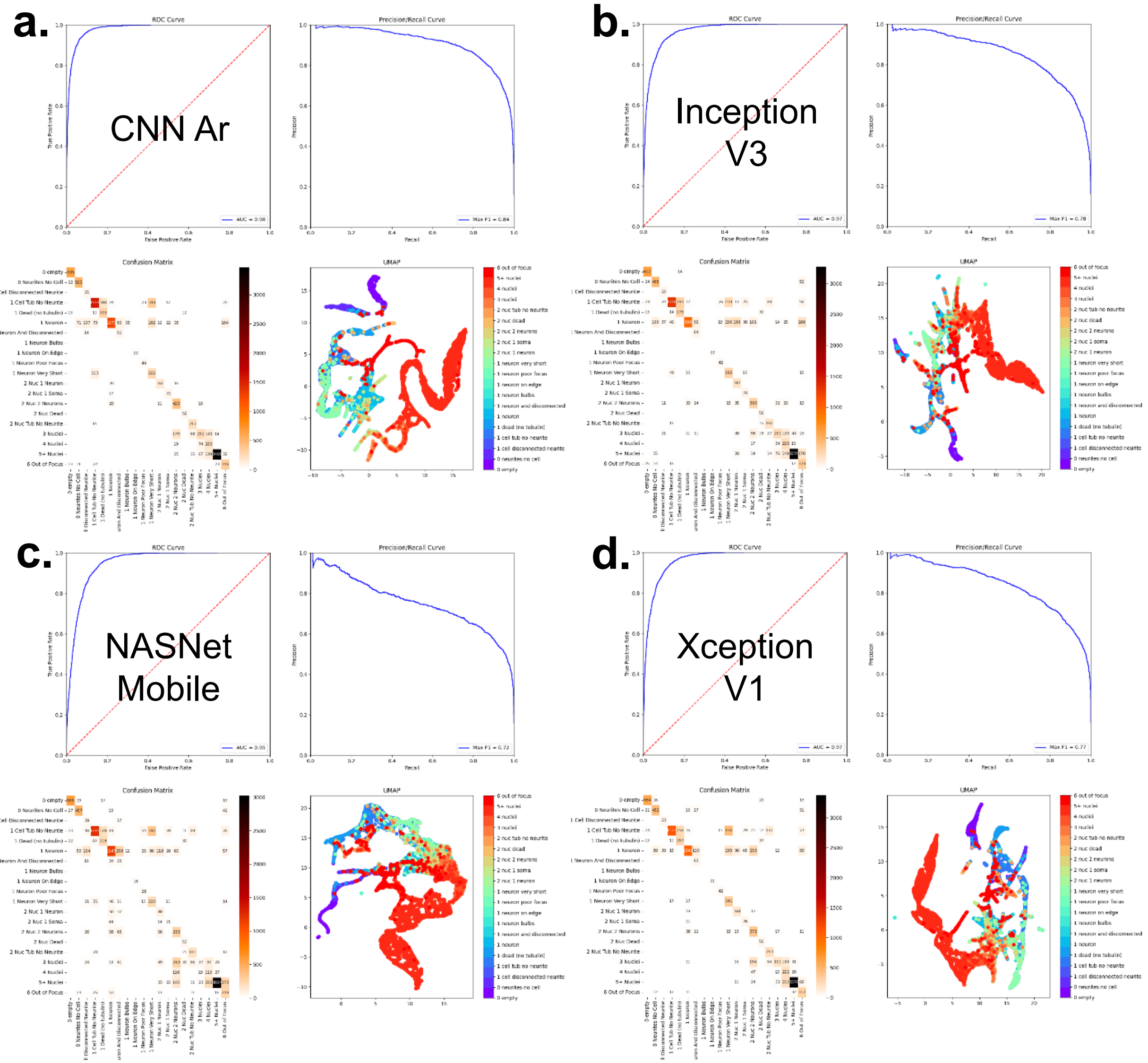


**Figure S8. Detail plots for non-CNN models.** Each panel shows an AUC ROC curve, F1 score (precision/recall) curve, confusion matrix for all 20 classes and a corresponding UMAP for a. CNN Ar, b. Inception V3, c. NASNet Mobile, and d. Xception V1 models.

a.

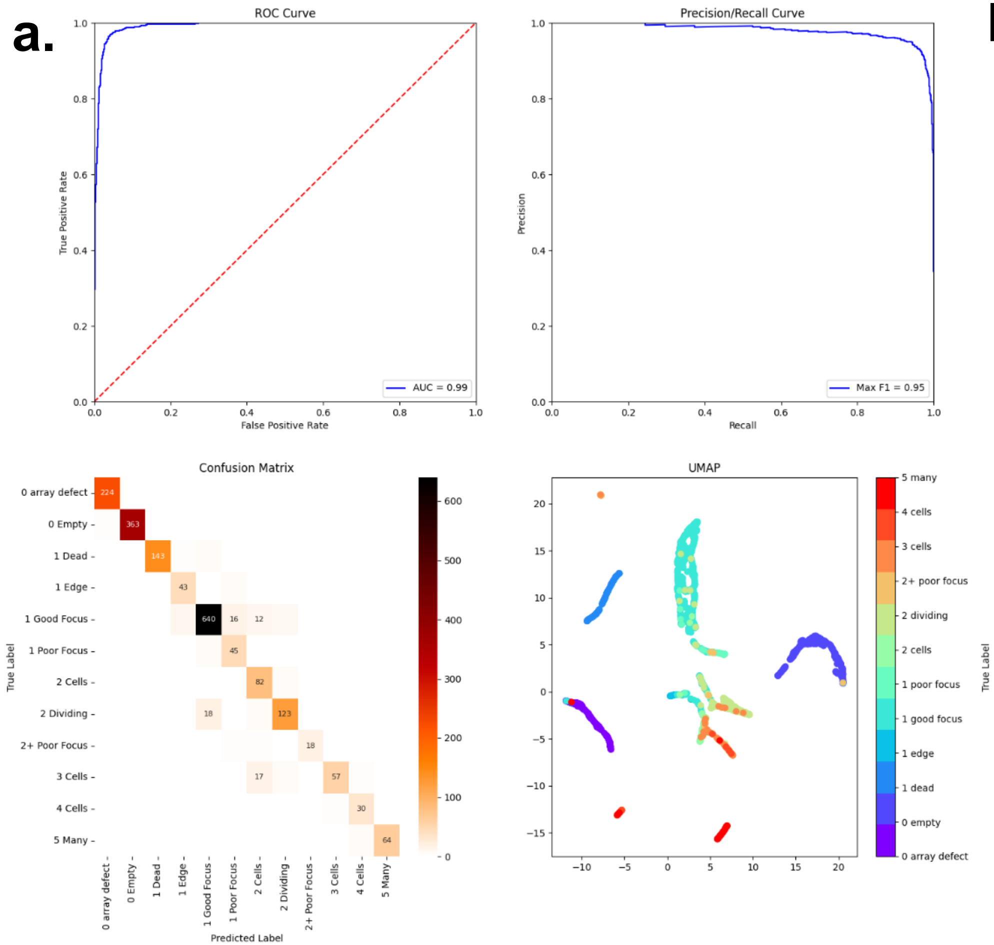


acc_0.8908 CNN_Nr CNNs_7 kn_4 flt_36 iRt_1.75
ds__194, 119_ nE_233,660 bSz_64 bNm_1 i_1

b.

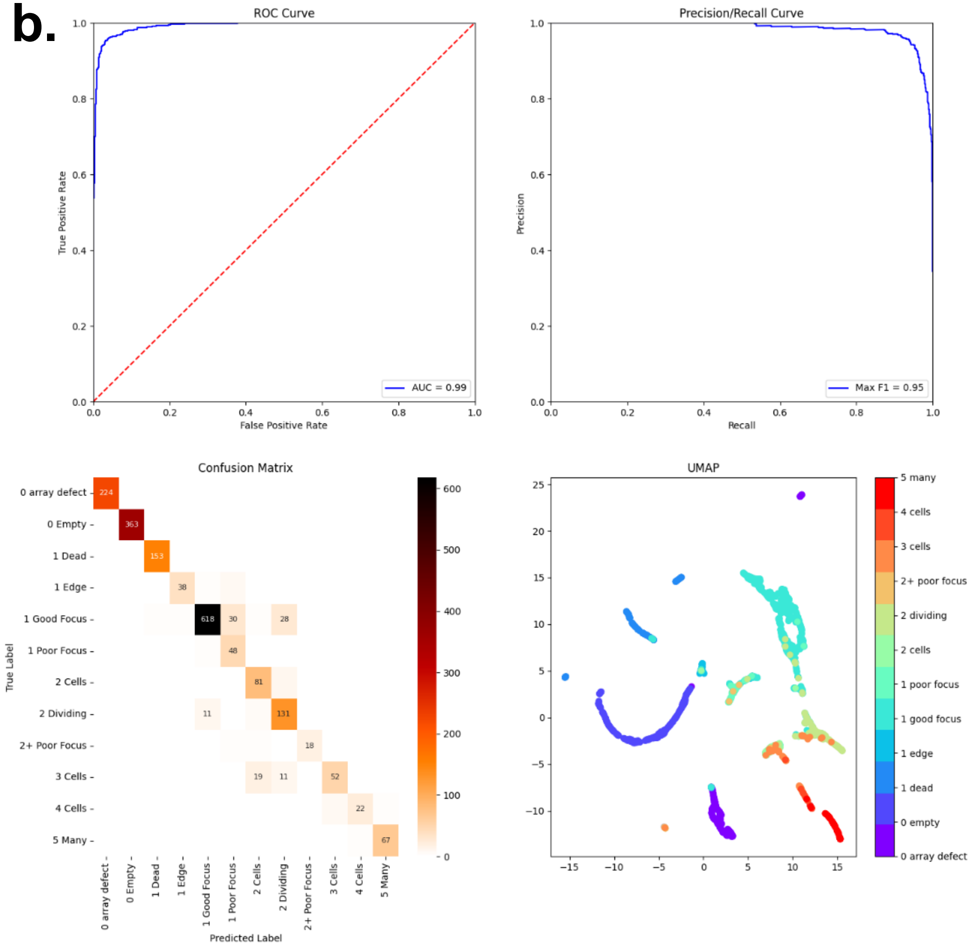


acc_0.8601 CNN_Nr CNNs_5 kn_4 flt_30 iRt_1.75
ds__703, 208, 108_ nE_370,660 bSz_64 bNm_0 i_53

**Figure S9. Detail plots for top 2 models.** Each panel shows an AUC ROC curve, F1 score (precision/recall) curve, confusion matrix for all 20 classes and a corresponding UMAP for a. the best and b. the second-best U2OS model.
